# Targeting protein tyrosine phosphatase non-receptor type 2 with a novel inhibitor for AML therapy

**DOI:** 10.1101/2024.05.07.592940

**Authors:** Wenbin Kuang, Jinxin Jiang, Xiao Wang, Dawei Wang, Minghui Ji, Yasheng Zhu, Kai Yuan, Jiayu Ding, Wenmu Wang, Mingge Song, Wenjian Min, Fei Huang, Liping Wang, Wanjian Gu, Haiping Hao, Yibei Xiao, Peng Yang

**Author notes:** Wenbin Kuang, Jinxin Jiang, Xiao Wang and Dawei Wang contributed equally to this article. Corresponding authors: Peng Yang,; #639 Longmian Avenue, Jiangning District, Nanjing, 211198, P. R. China. Yibei Xiao,; Haiping Hao,; Wanjian Gu,.

## Abstract

Acute myeloid leukemia (AML) is a fatal disease characterized by a bleak prognosis. For over four decades, treatment options for AML have been constrained to administering high-dose cytotoxic chemotherapy. However, the emergence of drug resistance and the resultant toxic side effects have created an urgent necessity for identifying novel therapeutic targets. In this study, non-receptor protein tyrosine phosphatase type 2 (PTPN2) is highly expressed in AML. Remarkably, knock down of PTPN2 expression markedly mitigated AML burden both in vitro and in vivo. Additionally, we unveiled the direct interaction between PTPN2 and C-MYC, establishing C-MYC as a direct acting substrate of PTPN2. Then, we performed a small-molecule compound screening to identify K73, a selective inhibitor of PTPN2. At the same time, K73 showed a good safety profile and exhibited strong activity against AML in vivo. In conclusion, we have pinpointed a considerable therapeutic potential in targeting PTPN2 for AML treatment and discovered a novel class of selective PTPN2 inhibitors suitable for AML therapy.

## Introduction

Acute myeloid leukemia (AML) is a complex and diverse disease characterized by the abnormal growth of myeloid lineage cells in the hematopoietic system^1-3^. In recent times, despite significant advancements in comprehension of AML’s pathogenesis and prognosis, the majority of patients diagnosed with AML still face restricted treatment alternatives^4,5^. Therefore, there is a need to develop new and effective anti-AML treatments.

PTPN2 is an important member of protein tyrosine phosphatases (PTPs)^6–12^, it has been identified as a potential target for tumor therapy^13–17^. PTPN2 shares a 72% genetic similarity and structural resemblance with PTP1B^18,19^. Despite the close relationship, PTP1B and PTPN2 demonstrate exceptional specificity in selecting their substrates^20–27^. PTPN2 plays a negative role in T-cell receptor (TCR) signaling by dephosphorylating and disabling LCK^28,29^. Additionally, PTPN2 also acts as an antagonist to cytokine signaling that is necessary for T-cell function, homeostasis, and differentiation^30–32^. PTPN2 deficiencies have been shown to promote PTK and STAT-3/5 signaling and tumorigenicity in cell lines, xenografts, and genetically engineered mouse models of cancer^33–36^. However, recent research has shown that PTPN2 deletion in melanoma and colon tumors promotes IFN/JAK-1/STAT-1 signaling, T cell recruitment, and antigen presentation, leading to improved anti-tumor immunity^13,37,38^. Thus, PTPN2’s role in cancer is complex and context-dependent, its deletion can enhance anti-tumor immune responses and inhibit tumor growth. In our previous studies, we demonstrated that PTPN2 is a key prognostic factor of pancreatic adenocarcinoma, and with pan-cancer analysis, we also found that PTPN2 is highly expressed in AML^39^. Nevertheless, the precise mechanisms by which heightened PTPN2 levels contribute to the development of AML remain elusive.

In this study, we investigated the profound impacts and underlying mechanisms of inhibiting PTPN2 on disrupting the cancer cell cycle. By conducting a variety of screening and validation experiments, we unraveled that C-MYC is a direct acting substrate of PTPN2 and further explored the therapeutic potential of targeting PTPN2 for the treatment of AML. Moreover, we have obtained a novel selective small molecule inhibitor of PTPN2 through a small-molecule screening platform, amplifying the wide-ranging possibilities in AML therapy.

## Results

### Increased PTPN2 expression in AML is associated with poor prognosis in AML patients

To investigate the role of PTPN2 in AML, The Cancer Genome Atlas (TCGA) was mined to analyze the expression of PTPN2 in normal and AML patients. We found the higher expression of PTPN2 in AML patients when compared with normal controls (Figure. 1A). We also found that the lower expression of PTP1B in AML patients when compared with normal controls (Supplementary Figure 1A). Furthermore, we analyzed the expression of PTPN2 gene in GSE30029 and GSE128103 chips, and the results showed that the expression of PTPN2 in AML samples were significantly higher than that in normal samples (Figure 1B and 1C). Then, as shown in Figure 1D, the PTPN2 mRNA levels increased, which were significantly correlated with the overall survival (OS) (p < 0.05). Meanwhile, we divided the bone marrow blood microarray sequencing results of 203 AML patients in the TARGET database into high expression group and low expression group according to the PTP1B expression value, and the overall survival rate was analyzed, revealing a significantly better prognosis for the high expression group (p < 0.01) (Supplementary Figure 1B). Based on the findings from the TARGET database, the expression of PTPN2 was categorized into three groups: high expression, medium expression, and low expression, subsequently, the overall survival rate was analyzed, the results revealed significant differences among these groups, indicating that the prognosis was notably worse in the high PTPN2 expression group (Figure. 1E). After, we analyzed Receiver Operating Characteristic (ROC) curves to explore the effect of PTPN2 expression on the deterioration of AML, the area under the curve (AUC) values of the ROC curves of PTPN2 were 0.96, which predicted that the expressions of PTPN2 might be related to the progression of AML (Figure. 1F). Then, the results of RT-qPCR and Western blot (WB) further showed that the mRNA and protein levels of PTPN2 in six blood samples from AML patients were higher than those in normal samples (Figure. 1G and 1H), and in AML cell lines, the protein level of PTPN2 in NB4 and MOLM13 cells was significantly higher than that in normal cells (Figure 1I). In conclusion, PTPN2 is highly expressed in both mRNA and protein levels in AML patient samples and AML cell lines, which is a potential drug target for the treatment of AML.

**Figure 1.**
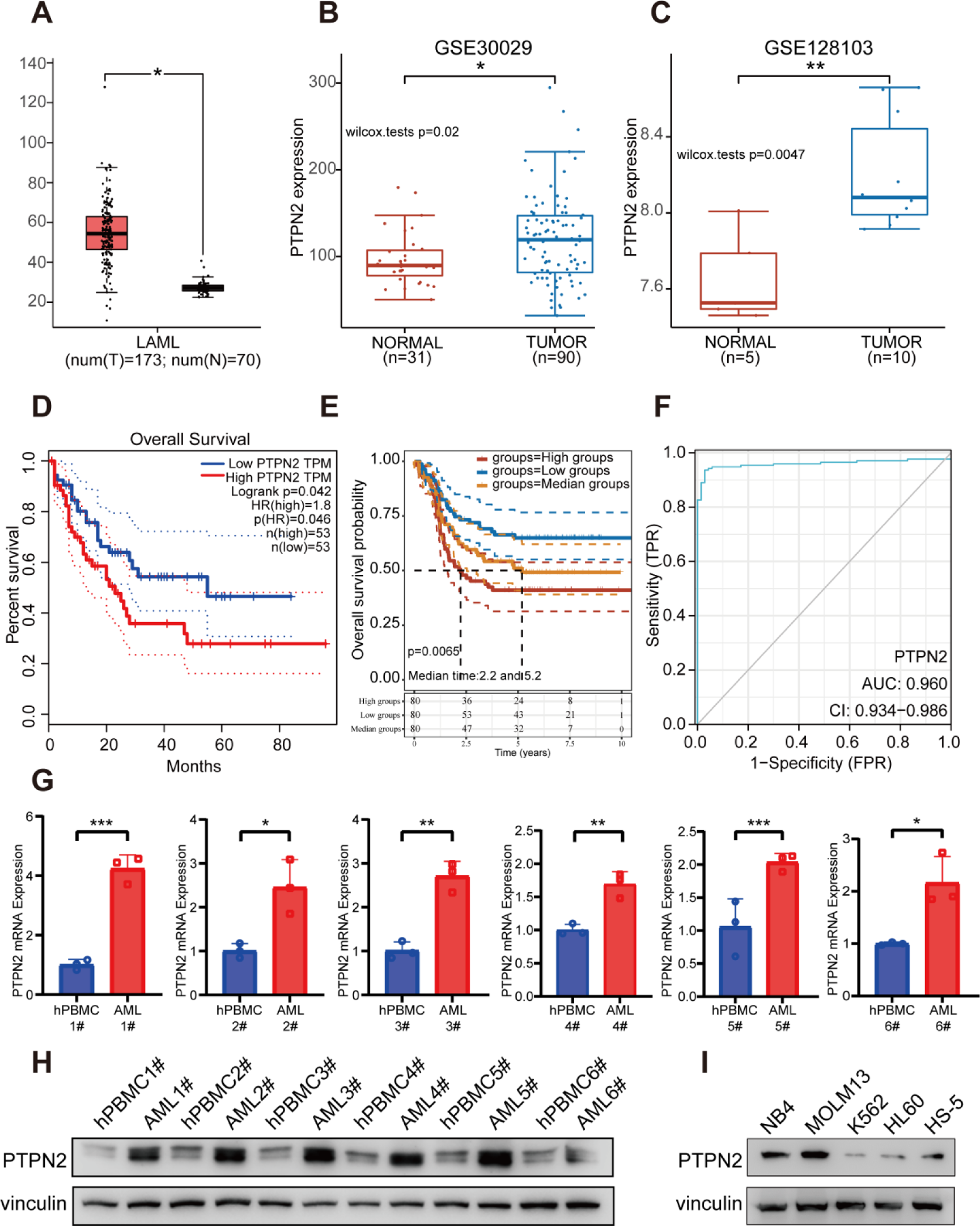
PTPN2 is highly expressed in AML and associated with poor survival. (A) Comparison of PTPN2 expression between tumors (red, n = 173) and normal (gray, n = 70) (B) PTPN2 expression analysis of monocytes from 31 normal samples and 90 tumor samples from GSE30029 chip (C) PTPN2 expression in the sequencing results of 5 normal umbilical cord blood cell samples and 10 tumor samples’ leukemic blasts, leukemic stem cells and lymphocytes samples from GSE128103 chip. The quantile percentile is illustrated by the whiskers of the box plot, where minima, 25%, median, 75%, and maxima are displayed from bottom to top, respectively. The application of a two-tailed Student’s t-test was conducted without considering adjustments for multiple comparisons, specifically the false discovery rate (FDR) (D) Kaplan-Meier survival plot of high (red line, n = 53) and low (blue line, n = 53) PTPN2 expression AML patients (data from TCGA). Statistical analysis was performed using log-rank test, P = 0.042 (E) Kaplan-Meier survival plot of high (red line), median (yellow line), and low (blue line) PTPN2 expression AML patients (data from the TARGET database). Statistical analysis was performed using log-rank test, P = 0.0065 (F) The receiver operating characteristic curve (ROC) predicts the accuracy of PTPN2 as a diagnostic factor for AML (G) PTPN2 mRNA level in tumor and hPBMCs in six AML patients. Data represent the mean ± SD. n = 3 biological replicates. Statistical analysis was performed using a two-tailed unpaired Student’s t-test (H) PTPN2 protein level in tumor and hPBMCs in six AML patients (I) PTPN2 protein levels of Human bone marrow stromal cell line HS-5 and four different Human myeloid leukemia cell lines. *, P < 0.05; **, P < 0.01; ***, P < 0.001.

### Knock down of PTPN2 significantly regulates the progression of AML

In order to explore regulation of PTPN2 on AML, PTPN2 was knocked down and over-expressed in NB4 and MOLM13 cells (Figure 2A, 2B and Supplementary Figure 2A). The growth capacity of NB4 and MOLM13 cells with PTPN2 knock down was significantly lower than that of NB4 and MOLM13 cells transfected with no loaded virus, while the over-expression of PTPN2 could rescue cell growth capacity (Figure 2A and 2B). The cell cycle results showed that NB4 and MOLM13 cells with PTPN2 knock down were arrested in G1 phase, conversely, over-expression of PTPN2 resulted in the restoration of cell cycle (Figure 2C and 2E). Similarly, knock down of PTPN2 could significantly increase apoptosis of NB4 and MOLM13 cells, and over-expression of PTPN2 resulted in the restoration of apoptosis (Figure 2D and 2F). Then, we expressed PTPN2 protein with mutation of amino acid 216 (Figure 2G) and found that PTPN2 can indeed be inactivated (Supplementary Figure 2B). Over-expressed the mutated PTPN2 on knock down NB4 and MOLM13 cells (Figure 2H and 2I), the growth capacity of NB4 and MOLM13 cells with knock down PTPN2 was significantly lower than that of cells transfected with no loaded virus, while the over-expression of the mutated PTPN2 could not rescue cell growth capacity (Figure 2H and 2I). Therefore, PTPN2 could regulate the cell proliferation, cell cycle, and apoptosis of NB4 and MOLM13 cells. Moreover, we subcutaneously implanted the MOLM13 shNC and shPTPN2 cells into the nude mice. Our findings demonstrated a lack of tumor formation in mice inoculated with MOLM13 cells that knocked down PTPN2, while mice inoculated with shNC cells exhibited noticeable tumor growth, and we observed no aberration in body weight among mice in comparison to the shNC group (Figure 2J).

**Figure 2.**
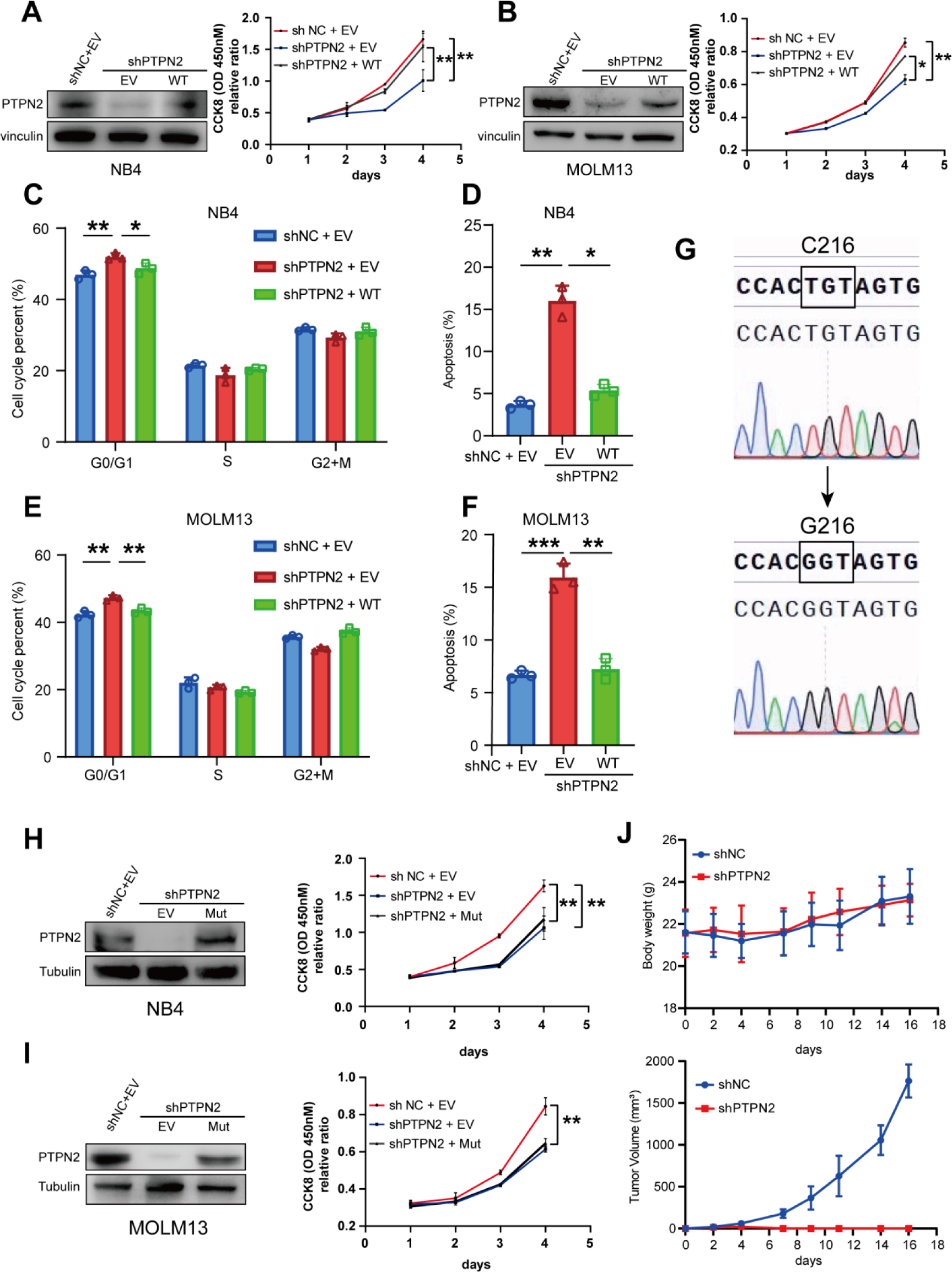
Depletion of PTPN2 inhibits AML in vitro and in vivo. (A) The protein level of PTPN2 in NB4 cells that were infected with lentiviral shNC, shPTPN2, and over-expression. Cell viability of NB4 shNC, shPTPN2, and over-expression cells during a 4-day course. Data represent the mean ± SD. n = 3 biological replicates. Statistical analysis was performed using a two-tailed unpaired Student’s t-test (B) Protein level of PTPN2 in MOLM13 cells that were infected with lentiviral shNC, shPTPN2, and over-expression. Cell viability of MOLM13 shNC, shPTPN2, and over-expression cells during a 4-day course. Data represent the mean ± SD. n = 3 biological replicates. Statistical analysis was performed using a two-tailed unpaired Student’s t-test (C and D) Cell cycle phase distribution and apoptotic rates of NB4 shNC, shPTPN2, and over-expression cells determined by flow cytometry. Data represent the mean ± SD. n = 3 biological replicates. Statistical analysis was performed using a two-tailed unpaired Student’s t-test (E and F) Cell cycle phase distribution and apoptotic rates of MOLM13 shNC, shPTPN2, and over-expression cells analyzed by flow cytometry. Data represent the mean ± SD. n = 3 biological replicates. Statistical analysis was performed using a two-tailed unpaired Student’s t-test (G) PTPN2 protein with mutation of amino acid 216, detected by Sanger sequencing (H and I) Protein level of PTPN2 in NB4 and MOLM13 shNC, shPTPN2, and over-expression PTPN2 C216G mutation (Mut) cells. Cell viability of cells during a 4-day course. Data represent the mean ± SD. n = 3 biological replicates. Statistical analysis was performed using a two-tailed unpaired Student’s t-test (J) Male BALB/c nude mice were implanted subcutaneously with MOLM13 shNC and shPTPN2 cells (n = 5 in each group). Tumor masses were photographed and weighed after executing mice. Body weights of mice were measured every two days. Data represent the mean ± SD. Statistical analysis was performed using a two-tailed unpaired Student’s t-test. *, P < 0.05; **, P < 0.01; ***, P < 0.001.

We also over-expressed PTPN2 in K562 cells (Supplementary Figure 2C), over-expression of PTPN2 in K562 cells significantly promoted the growth of K562 cells (Supplementary Figure 2D). Then, in order to further verify the regulatory role of PTPN2 in AML, we have separated AML primary cells from the other two blood samples from AML patients, and PTPN2 was knocked down in AML primary cells (Supplementary Figure 3A). The cell viability of AML primary cells was notably reduced when PTPN2 was knocked down, compared to AML primary cells transfected with an empty virus (Supplementary Figure 3B), Furthermore, the cell cycle analysis indicated that AML primary cells with PTPN2 knock down were predominantly arrested in the G1 phase (Supplementary Figure 3C), Additionally, the decrease in PTPN2 expression led to increase in apoptosis of AML primary cells (Supplementary Figure 3D). In summary, the significant decrease in AML burden both in vitro and in vivo was observed with the down-regulation of PTPN2, underscoring its vital role in the regulation of AML.

### Effect of PTPN2 knock down on MOLM13 cell signaling pathway

In order to analyze which genes and signaling pathways are involved in PTPN2 regulation of AML, transcriptome sequencing analysis (RNA-Seq) was performed on MOLM13 cells with PTPN2 knock down and MOLM13 cells transfected with no loaded virus. The results showed that 479 genes were significantly up-regulated and 527 genes were significantly down-regulated in the knock down PTPN2 group compared with the shNC group (Figure 3A). Then, the Hallmark enrichment analysis showed that down-regulated genes were mainly enriched in E2F targets, oxidative phosphorylation, G2M checkpoint, MYC target V1 and MYC target V2 pathways (Figure 3B), and up-regulated genes were mainly enriched in P53 pathway etc. (Figure 3C). Genome-wide enrichment analysis and GSEA analysis showed that PTPN2 knock down could significantly down-regulate MYC targets V1, MYC targets V2, G2M checkpoint, E2F targets and up-regulate P53 pathway, which suggested that knock down of PTPN2 is associated with inhibition of cell cycle and cell proliferation (Figure 3D and 3E). Then, WB assay showed that knock down of PTPN2 down-regulated CDK4, CDK6, P-Rb, C-MYC, BCL-2 and XIAP. At the same time, P21, P27 and P53 were up-regulated (Figure 3F). These results suggest that knock down of PTPN2 can inhibit cells through regulating cell cycle, apoptosis, and proliferation.

**Figure 3.**
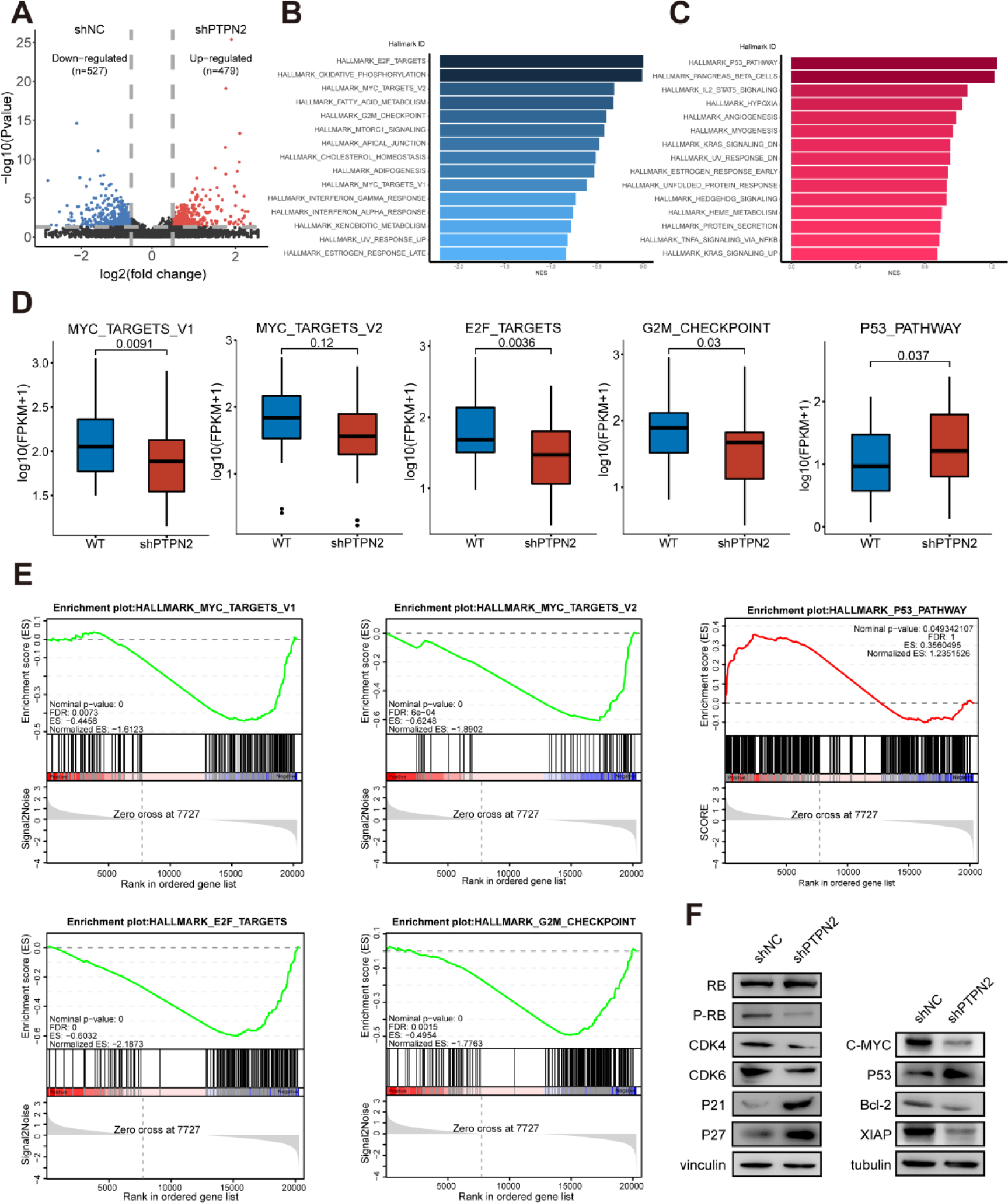
RNA Sequencing combined validation assays to identify affected signaling pathways of PTPN2 loss in AML cells. (A) Volcano plot of significantly affected (absolute fold change > 2, P value < 0.05) genes in MOLM13 shPTPN2 group relative to shNC group revealed by RNA-seq. P values were calculated using R v4.0.3 software (B and C) Bar graphs show the core-enriched down-regulated (blue) and up-regulated (red) signaling pathways in shPTPN2 groups when compared with shNC groups, respectively (D) The relative abundance of genes involved in MYC targets V1, MYC targets V2, E2F targets, G2M checkpoint and P53 pathway in MOLM13 PTPN2 KD cells. The quantile percentile is illustrated by the whiskers of the box plot, where minima, 25%, median, 75%, and maxima are displayed from bottom to top, respectively. The application of a two-tailed Student’s t-test was conducted without considering adjustments for multiple comparisons, specifically the false discovery rate (FDR) (E) The signaling pathways enriched in different groups obtained through Gene Set Enrichment Analysis (GSEA) (F) Immunoblotting analysis of protein on MOLM13 shPTPN2 and shNC cells. The experiments in F were repeated three times with similar results.

### C-MYC is a novel PTPN2 substrate

To investigate the molecular mechanism of PTPN2 function and investigate the potential target of PTPN2, high-throughput quantitative phosphoryl proteomics analysis was performed on MOLM13 cells with PTPN2 knocked down (Figure 4A). A total of 22,454 phosphoric acid sites were identified, of which 19,947 (88.83%) were serine, 2,309 (10.28%) were threonine, and 198 (0.88%) were tyrosine (Supplementary Figure 4A). Our comprehensive analysis of significantly up-regulated and down-regulated phosphorylated proteins revealed major pathways in which PTPN2 might be involved (Supplementary Figure 4B). We also analyzed the proteins binding to PTPN2 by immunoprecipitation mass spectrometry, and the results showed that 31 proteins could bind to PTPN2 (Supplementary Figure 4C and 4D), and C-MYC appeared in between up phosphorylated proteins and PTPN2-binding proteins (IP) (Figure 4B). Then, we focused on the target of C-MYC, which is involved in regulating cell proliferation, differentiation, growth, apoptosis, cell cycle progression, intracellular biomolecular metabolism, and malignant transformation of cells^40–42^. Heat maps showed that phosphorylation levels of P-C-MYC (S62) and P-C-MYC (T58) were up-regulated (Supplementary Figure 4E). The mass spectrograms of C-MYC proteins pT58 and pS62 are shown in Figure 4C and Supplementary Figure 4F. They were clearly fragmented and could be used for peptide recognition and site localization.

**Figure 4.**
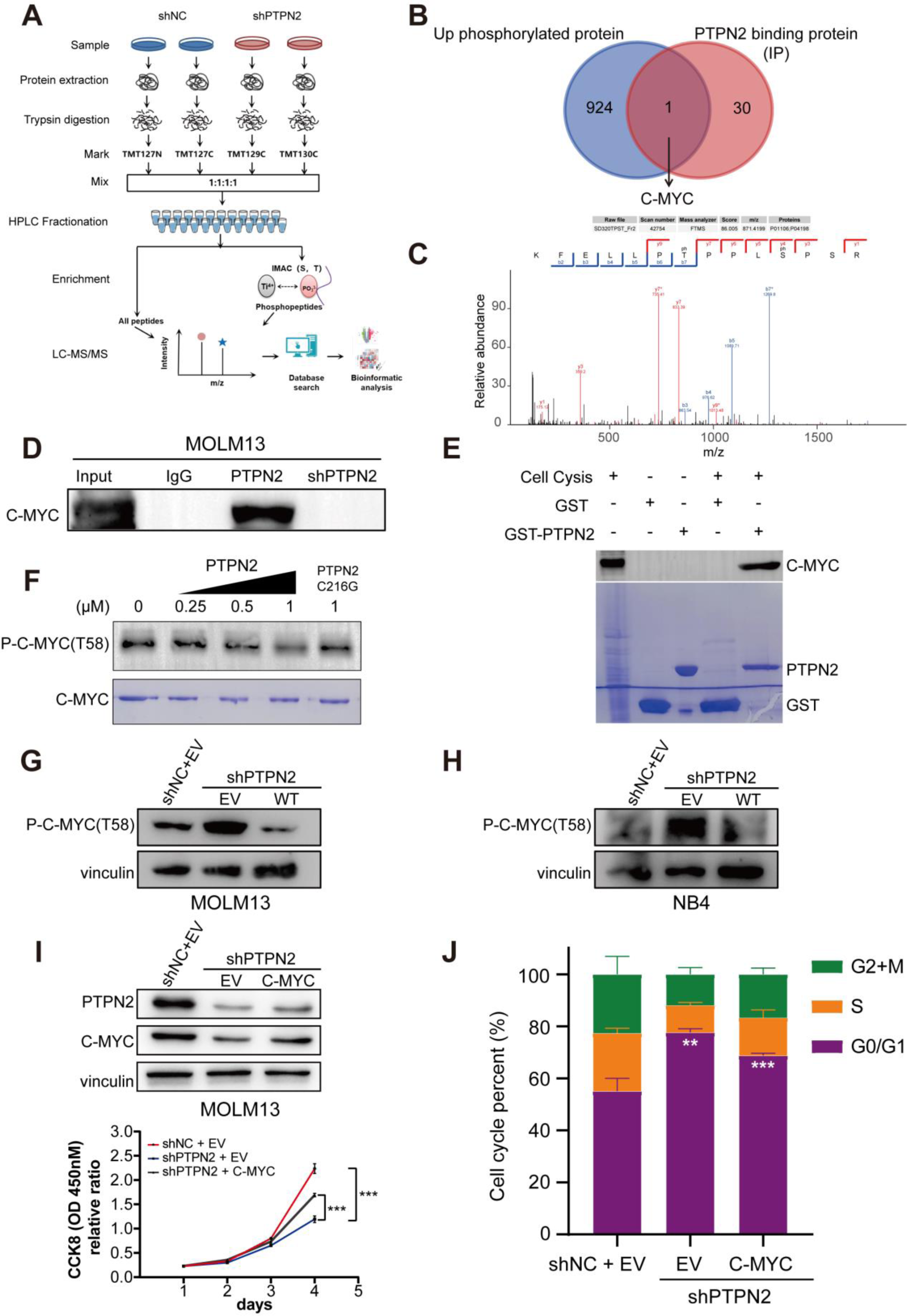
Quantitative phosphoproteomic analysis of MOLM13 cells with PTPN2 knock down and validation of PTPN2 substrates. (A) Workflow of the phosphoproteomic approach. samples were separately lysed, digested and TMT marked and the phosphorylated peptides were enriched by the IMAC tip and analyzed by LC-MS/MS (B) Venn diagrams show the overlap of up phosphorylated proteins and PTPN2-binding proteins (IP) (C) MS/MS spectra of the phosphosites of the potential PTPN2 substrates, pT58 of C-MYC (D) Validation of interaction of C-MYC with PTPN2 by co-immunoprecipitation assay (E) Validation of interaction of C-MYC with PTPN2 by GST pull assay (F) Dephosphorylated MYC T58 levels by PTPN2 were assessed by western blot in an in vitro dephosphorylation assay (G and H) Protein level of P-C-MYC (T58) in MOLM13 and NB4 cells that were infected with lentiviral shNC, shPTPN2 and over-expression (I) Protein level of PTPN2 and C-MYC in MOLM13 shNC, shPTPN2 and over-expression C-MYC cells. Cell viability of cells during a 4-day course. Data represent the mean ± SD. n = 3 biological replicates. Statistical analysis was performed using a two-tailed unpaired Student’s t-test (J) Cell cycle phase distribution of MOLM13 shNC, shPTPN2, and over-expression C-MYC cells analyzed by flow cytometry. Data represent the mean ± SD. n = 3 biological replicates. Statistical analysis was performed using a two-tailed unpaired Student’s t-test. **, P < 0.01; ***, P < 0.001.

Then, we detected the binding effect of PTPN2 and C-MYC using CO-IP assay, the WB results showed that PTPN2 could bind C-MYC in MOLM13 cells, but could not bind C-MYC when PTPN2 was knocked down (Figure 4D). The GST pull-down was used to detect the interaction between PTPN2 and C-MYC. WB results showed that the negative control GST label protein could not bind C-MYC in MOLM13 cell lysate, while the GST-PTPN2 recombinant protein could bind C-MYC in MOLM13 cell lysate (Figure 4E). These results fully confirmed that PTPN2 could interact with endogenous C-MYC. We also performed an in vitro phosphatase assay using purified PTPN2 and C-MYC proteins. As shown in Figure 4F, PTPN2 effectively dephosphorylated C-MYC T58, while the phosphatase-deficient PTPN2 mutant (C216G) was inactive. WB results showed that knock down of PTPN2 up-regulated the expression of P-C-MYC (T58) in MOLM13 and NB4 cells, while over-expression of PTPN2 down-regulated the expression of P-C-MYC (T58) in cells (Figure 4G and 4H). In summary, C-MYC is the direct substrate of PTPN2.

Then, we examined whether C-MYC would mediate PTPN2 function in AML cell lines. We over-expressed C-MYC in MOLM13 cells with PTPN2 knock down (Figure 4I). The growth ability of MOLM13 cells with PTPN2 knock down was significantly lower than that of MOLM13 cells transformed into empty virus, but over-expression of C-MYC could rescue cell growth capacity (Figure 4I). The results of cell cycle assay showed that MOLM13 cells with PTPN2 knock down were significantly arrested in G1 phase, and over-expression of C-MYC in MOLM13 cells with PTPN2 knock down could result in the restoration of cell cycle (Figure 4J).

In order to deeper analysis of potential PTPN2 target proteins and the effects of phosphoserine on MYC function, we performed large-scale genomic analyses of the TCGA database and found that PTPN2 deletion and MYC amplification or missense mutation were found to be almost mutually exclusive in AML (Supplementary Figure 5A). The reduction in C-MYC protein levels resulting from PTPN2 knock down can be reversed by the proteasome inhibitor MG132, suggesting that PTPN2 knock down influences the stability of C-MYC protein (Supplementary Figure 5B). The phosphorylation of C-MYC at T58 plays a crucial role in the degradation of C-MYC induced by PTPN2 knock down, as the C-MYC T58 mutant, which cannot be phosphorylated at this site, is not as readily degraded by PTPN2 knock down (Supplementary Figure 5C). We also over-expressed C-MYC and C-MYC T58 mutation protein in MOLM13 and NB4 cells with PTPN2 knock down (Supplementary Figure 5D and 5E), and the growth ability of MOLM13 and NB4 cells with PTPN2 knock down and cells over-expression of C-MYC T58 mutation protein was significantly lower than that of cells over-expression of C-MYC (Supplementary Figure 5D and 5E). The findings indicate that reducing PTPN2 levels enhances T58 phosphorylation of C-MYC, leading to degradation of C-MYC.

### The development of a selective PTPN2 inhibitor K73 significantly inhibited growth of AML cells in vitro

In order to identify PTPN2 inhibitors, we conducted in vitro activity detection of PTPN2 and screened 7769 compounds from various libraries (including the Selleck compound library, FDA drug library, natural active product library, and our house library) using high throughput methods (Figure 5A). We found the compound NP-12 which had good inhibitory activity against PTPN2 phosphatase activity, and its inhibitory rate of PTPN2 phosphatase activity is 99% at 10 μM concentration (Supplementary Figure 6A). The IC_50_ value of NP-12 for PTPN2 was 3.23 μM and for PTP1B was 4.66 μM (Supplementary Figure 6B and 6C). The binding pattern of NP-12 to PTPN2 indicates that oxygen on 1,2, 4-thiadiazolidin-3,5-dione forms a hydrogen bond interaction with GLN-260 (Supplementary Figure 6D). Then, through systematic optimization (see Methods for details), multiple series of derivatives K72-K82 (Supplementary Figure 7) were synthesized. Among them, compound K73 exhibited the most potent PTPN2 inhibitory activity with an IC_50_ value of 323.6 nM (Figure 5B), while the IC_50_ value of compound K73 against PTP1B was 4.27 μM (Figure 5C). The binding pattern of the compound K73 with PTPN2 showed that the oxygen atoms on 1,2,4-thiadiazolidin-3,5-dione formed hydrogen bond interaction with GLN-260, and the oxygen atoms and nitro atoms on 4-nitrophenoxy group formed hydrogen bond interaction with TRY-48 and CYS-216 (Supplementary Figure 6E). In the MST PTPN2 interaction assay, titration of compound K73 in labeled PTPN2 protein resulted in a thermophoresis shift with a K_d_ of 646 nM (Figure 5D) and titration of compound K73 in labeled PTPN2 mutation protein resulted in a thermophoresis shift with a K_d_ of 31.57 μM (Supplementary Figure 6F). We also used SPR to tested compound K73 bind to PTPN2, PTPN2 has a clear concentration-dependent response to compound K73, with a calculated K_d_ of 3.96 μM (Supplementary Figure 6G), it suggests that the compound K73 binds closely to PTPN2.

**Figure 5.**
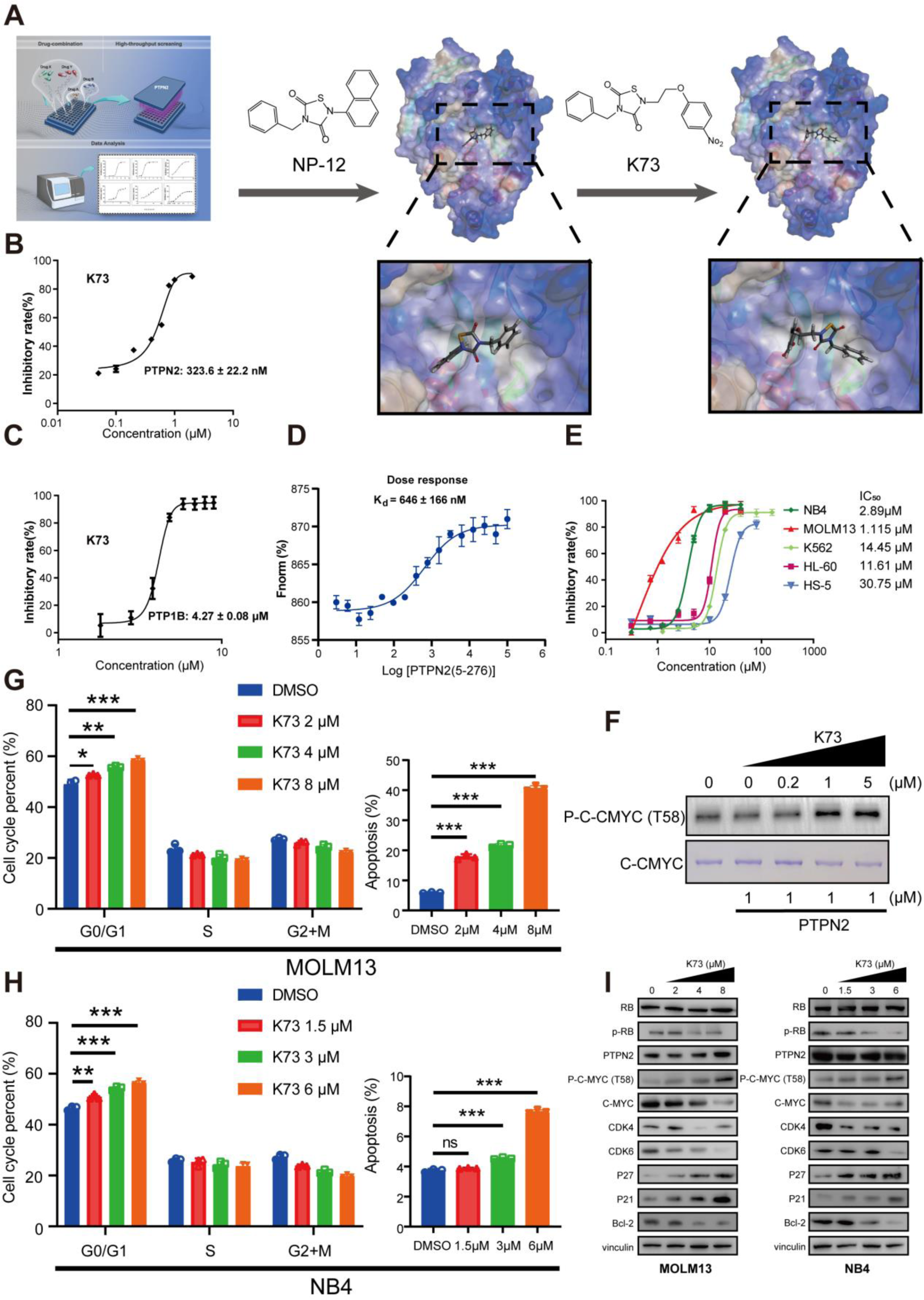
Discovery of selective PTPN2 inhibitor K73 and K73 inhibited cell growth, arrest cell cycle, and induced apoptosis in AML cells. (A) High-throughput screening flow diagram for the discovery of the PTPN2 inhibitor compound NP-12, and chemical structure of K73 (B and C) Inhibitory activity of compound K73 against PTPN2 and PTP1B. Data represent the mean ± SD. n = 3 biological replicates. Statistical analysis was performed using a two-tailed unpaired Student’s t-test (D) MST curve for interaction of K73 with PTPN2 and employing standard data analysis with Mo. Affinity Analysis v2.2.4 software. Data represent the mean ± SD. n = 3 biological replicates. Statistical analysis was performed using a two-tailed unpaired Student’s t-test (E) K73 inhibiting the proliferation of a variety of AML cell lines. Data represent the mean ± SD. n = 3 biological replicates. Statistical analysis was performed using a two-tailed unpaired Student’s t-test (F) Dephosphorylated MYC T58 levels by K73 inhibit PTPN2 were assessed by western blot in vitro dephosphorylation assay (G) Cell cycle phase distribution and apoptotic rates of MOLM13 cells s after treatment with DMSO, K73 (2, 4, 8 µM) for 24 h determined by flow cytometry. Data represent the mean ± SD. n = 3 biological replicates. Statistical analysis was performed using a two-tailed unpaired Student’s t-test (H) Cell cycle phase distribution and apoptotic rates of NB4 cells after treatment with DMSO, K73 (1.5, 3, 6 µM) for 24 h determined by flow cytometry. Data represent the mean ± SD. n = 3 biological replicates. Statistical analysis was performed using a two-tailed unpaired Student’s t-test (I) The total proteins in MOLM13 and NB4 cells were extracted and used in the western blotting analysis of indicated proteins. *, P < 0.05; **, P < 0.01; ***, P < 0.001; ns, not significant.

We evaluated the inhibition of compound K73 on the proliferation of AML cells by Cell Counting KIT-8 (CCK8) assay. Compound K73 inhibited MOLM13 and NB4 cells with high expression of PTPN2 with IC_50_ values of 1.11 μM and 2.89 μM, respectively. However, K562 and HL-60 cells with low expression of PTPN2 had poor inhibitory activity, IC_50_ values were 14.45 μM and 11.61 μM, respectively, which showed no significant toxicity to HS-5 cells, IC_50_ value was 30.75 μM (Figure 5E). However, compound K73 inhibited MOLM13 shPTPN2 cells with IC_50_ values greater than 40 μM (Supplementary Figure 6H). These results indicated that compound K73 could significantly inhibit the proliferation of AML cells, and the inhibitory effect was more obvious on AML cells with high PTPN2 expression, indicating that K73 had good target selectivity. Compound K73 significantly inhibited the growth of MOLM13 and NB4 cells in a concentration-dependent manner (Supplementary Figure 6I and 6J). To further investigate the mechanism of action, compound K73 was incubated with purified PTPN2 protein in vitro, and then phosphatase assay was performed with C-MYC protein in vitro. The results showed that compound K73 inhibited PTPN2 and promoted C-MYC T58 phosphorylation in a dose-dependent manner, suggesting that compound K73 could inhibit PTPN2 activity in vitro (Figure 5F). In addition, compound K73 could arrest the cell cycle of MOLM13 and NB4 cells in G1 phase in a concentration-dependent manner (Figure 5G and 5H) and promote the apoptosis of MOLM13 and NB4 cells in a concentration-dependent manner (Figure 5G and 5H). Then, WB assay showed that compound K73 could significantly up-regulate the expression of P-C-MYC (T58), P21 and P27, C-MYC, CDK4, CDK6, P-Rb and Bcl-2 were significantly down-regulated in MOLM13 cells and NB4 cells (Figure 5I). Then, the Jurkat cell line was chosen as the cell model for investigating the effects of compound K73. The Jurkat cell line is a well-established immortalized cell line of human T lymphocytes that is commonly employed in studies related to T cell signaling. After subjecting the Jurkat cell line to a 24-hour treatment with compound K73, we observed a notable increase in STAT1 phosphorylation. This finding suggests that compound K73 has the ability to activate the downstream signaling pathways by inhibited PTPN2 in the Jurkat cells (Supplementary Figure 6K). In summary, knock down of PTPN2 and the use of the small-molecule inhibitor K73 had similar effects on AML cells when tested in vitro. Both methods inhibited cell growth by causing G0/G1 arrest and inducing apoptosis.

### Evaluation of anti-AML activity of compound K73 in vivo

In order to evaluate the safety of compound K73 in vivo. ICR mice were divided into four groups. In the three groups, the doses were 1000mg/kg, 2500 mg/kg, and 5000 mg/kg, respectively, and one control group, which were given normal saline. After 14 days of observation, the mice in each group did not die, and no abnormalities were observed. There was no significant difference in body weight between the three groups and the control group (Figure 6A). Heart, liver, spleen, lung, and kidney weights of mice in the three dosing groups did not differ significantly from those in the control group (Figure 6B). H&E staining of Heart, liver, spleen, lung, and kidney tissues indicated that compound K73 exhibited a good safety profile in vivo (Figure 6C).

**Figure 6.**
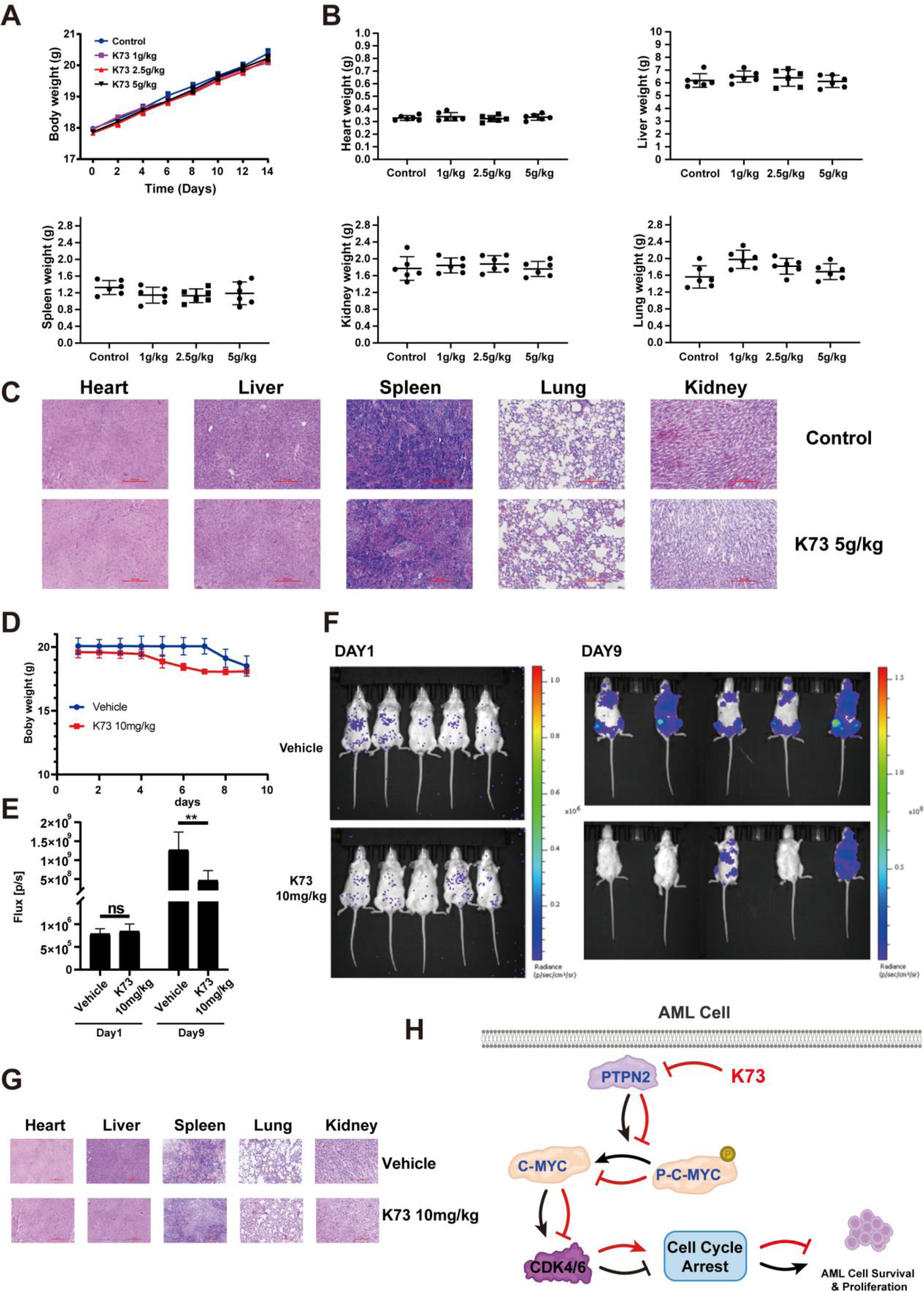
Compound K73 exhibits a good anti-AML effect in vivo. Mice were given vehicle or K73 (1000 mg/kg, 2500 mg/kg, and 5000 mg/kg) orally. (A) Following the administration of a single dosage, the mice’s body weight was assessed every two days. Notably, over a period of 14 days, no mortalities among the mice were observed, prompting the execution and subsequent dissection of all subjects (B) The vital organs of mice, comprising the heart, liver, spleen, lung, and kidney, were removed and subjected to weighing. Error bars, mean ± SD, n = 10 (C) H&E staining of main organs from mice treated with vehicle or K73. D-G To establish the MOLM13-Luciferase transplantation tumor model, M-NSG mice were injected with 5 × 10^5^ tumor cells through the tail vein. After 2 days of inoculation, the mice with tumors were divided randomly into two groups (n = 5/group) based on the Flux value. One group received the vehicle every day, while the other group was administered K73 at a dosage of 10 mg/kg daily (D) body weight of mice was measured every 2 days. Error bars, mean ± SD, n = 5 (E and F) Small animal imaging to detect the fluorescence intensity of mice on day 2 and day 9 (G) H&E staining of main organs from mice treated with vehicle or K73. Error bars, mean ± SD, n = 5 (H) Proposed working model for K73 inhibits PTPN2 in the treatment of AML. **, P < 0.01.

Then, we established an animal model by injecting Molm13-luciferase cells into the tail vein of M-NSG mice. The experimental animals were divided into 2 groups: control group and compound K73 group (10 mg/kg). Both in the control group and the drug administration group, the weight of mice decreased significantly on the 9th day, and the death occurred in the control group on the 10th day (Figure 6D). We used small animal imaging to detect the fluorescence intensity of mice on day 2 and day 9, as shown in Figure 6E and 6F, the fluorescence intensity of mice in the drug group was significantly lower than that in the control group on day 9. The H&E staining of the heart, liver, spleen, lung, and kidney tissues of the two groups of mice did not show obvious toxicity (Figure 6G). This result indicated that compound K73 had a good anti-AML effect.

We inferred the mechanism of action of compound K73, as shown in Figure 6H, compound K73 promoted the phosphorylation of C-MYC T58 by inhibiting the activity of PTPN2, thereby increasing the proteasome-mediated degradation of C-MYC, leading to the down-regulation of CDK4/6, which arrested the cell cycle in G1 phase, and finally achieved the effect of AML treatment.

## Discussion

Acute myeloid leukemia (AML) is a complex and life-threatening hematological condition that impacts the blood, bone marrow, and lymphoid systems of the body^43,44^, As our comprehension of the molecular genetic changes underlying the pathogenesis of AML has expanded, recent research has primarily focused on addressing intracellular pathways regulated by these anomalous proteins. The potential of molecular targeting as a prospective approach to treating AML is significant^45–48^.

There have been remarkable developments in the progress of drugs that target PTPNs family, these medications have proven effective in treating diverse human conditions^49,50^. PTPN2 plays a key role within the PTPNs family, with prior research indicating that deactivation of PTPN2 may have a role in the formation of tumors by enhancing PTK signaling^33–36^, however, it cannot be ruled out that the removal of PTPN2 within the framework of other cancer-causing incidents^51^. The response of IFNγ/JAK/STAT-1 is of utmost importance in the progression of immunotherapy in various tumors^52,53^, furthermore, it plays a crucial role in anti-cancer immunity and cellular therapies which utilize T cells/CAR-T cells^52,54,55^. Recent studies have shown that PTPN2 deletion promotes IFN/JAK-1/STAT-1 signaling, T cell recruitment, and antigen presentation in melanomas and colon tumors, thereby enhancing anti-tumor immunity^13,37,38^. In our prior investigations, we substantiated that PTPN2 is a key prognostic factor of pancreatic adenocarcinoma, in addition, pan-cancer analysis showed that PTPN2 is highly expressed in AML^39^. Thus, the role of PTPN2 in cancer is complex and context-dependent. Taking into consideration the crucial role of PTPN2 in cancers, we conducted data mining utilizing TCGA and gene chip approaches. The analysis revealed that PTPN2 exhibited high expression levels in AML, which displayed a positive correlation with both clinical prognosis and mortality (Figure 1A-1E). Furthermore, we observed a heightened expression of PTPN2 in both AML patient samples and cells (Figure 1G-1I).

To further verify the regulatory effect of PTPN2 on AML cells, genetic inhibition of PTPN2 significantly reduce the burden of AML (Figure 2A-2F). In addition, knock down of PTPN2 cells resulted in their loss of tumorigenicity in mice (Figure 2J). These findings collectively suggest that PTPN2 represents a promising therapeutic target for combating AML. Then, we conducted transcriptome sequencing analysis (RNA-Seq) on MOLM13 cells. The Hallmark enrichment analysis results showed that down-regulated genes were mainly enriched in E2F targets pathway etc, which suggested that knock down of PTPN2 is associated with inhibition of cell cycle and cell proliferation (Figure 3B and 3C). Then, high-throughput protein phosphorylation and omics analysis identified C-MYC as a direct substrate of PTPN2 (Figure 4A-4J). Therefore, PTPN2 has significant potential as a target for the development of drugs to effectively address AML.

According to TCGA database analysis, the high expression of PTPN2 in AML is positively correlated with clinical prognosis, while PTP1B is opposite (Supplementary Figure 1), the development of selective PTPN2 small molecule inhibitors is beneficial for the treatment of AML. Then, we found that NP-12 had a good inhibitory activity against PTPN2 through high-throughput screening of compounds in the compound library (Supplementary Figure 6A). Furthermore, a new derivative K73 was obtained by initial structural optimization of NP-12, which showed good inhibitory activity against PTPN2 with an IC_50_ value of 323.6 nM (Figure 5B), while the IC_50_ value of compound K73 against PTP1B was 4.27 μM (Figure 5C), indicating that compound K73 had selectivity for PTPN2. The results of in vitro experiments showed that compound K73 had a stronger inhibitory effect on the proliferation of AML cells with high PTPN2 expression than that of AML cells with low PTPN2 expression and could inhibit PTPN2 activity in vitro, and then block the cell cycle in G1 phase in the cell, promoting the apoptosis of AML (Figure 5E-5H). And compound K73 has shown a good anti-AML effect in vivo (Figure 6E). C-MYC, when phosphorylated at T58, is identified by the E3 ubiquitin ligase FBXW7 and subsequently undergoes degradation via the 26S proteasome^56–59^. These results suggest that compound K73 facilitated the phosphorylation of C-MYC T58 by suppressing the function of PTPN2. this led to the enhancement of C-MYC’s proteasomal degradation, ultimately resulting in the down-regulation of CDK4/6 and caused the arrest of the cell cycle in the G1 phase, thereby achieving the desired therapeutic outcome for AML (Figure 6H).

In summary, our findings reveal a previously unrecognized activity of PTPN2 and its mechanism of action on AML regulation. Additionally, we obtained a selective PTPN2 inhibitor, which could serve as a small-molecule probe for further biological studies. Furthermore, since PTPN2 is involved in critical processes in various types of human cancers, targeting this protein could potentially benefit patients with other refractory cancers as well.

## Methods

### Molecular Cloning

To obtain the recombinant proteins PTPN2 (1-314 AA) and PTP1B (1-321 AA). We first extracted RNA from HEK293T cells and obtained cDNA by reverse transcription reaction, and then used cDNA as a template to obtain the target genes PTPN2 and PTP1B by polymerase chain reaction (PCR) using specific primers. We used homologous recombination to homologize the target genes PTPN2, PTP1B, and the vector pGEX6p-1 by reverse complementary pairing of homology arms. The in vitro recombination product was transformed into clone strain DH5α, verified by colony PCR, and then sequenced using Sanger gene sequencing, and the correctly sequenced single clone was extracted into plasmid and transformed into expression strain BL21 (DE3).

For the mutant PTPN2 C216G (1-314 AA), we designed point mutation primers, amplified the plasmid fragments by rolled-loop PCR and verified the location of the plasmid fragments by agarose gel electrophoresis, transfected DH5α after DpnI digestion, picked a single clone to be sequenced to verify the mutation site, and extracted the plasmid for transformation of BL21 (DE3) after correct sequencing.

In order to overexpress C-MYC (1-415 AA) at the cellular level, we used cDNA from HEK293T cells as a template and PCR to obtain the target gene C-MYC, which was homologously recombined by homologous recombination with pCDNA3.1(+) vector, and plasmid extraction was performed after correct sequencing. Similarly, we used point mutation to mutate threonine 58 of C-MYC (1-415 AA).

Mutation primers:

F-PTPN2(C216G): ATCCACGGTAGTGCAGGCATTGGGCG

R-PTPN2(C216G): TGCACTACCGTGGATCACCGCAGGCC

F-C-MYC(T58A): TGCTGCCCACCCCGCCCCTGTCCCCTAGCC

R-C-MYC(T58A): GCGGGGTGGGCAGCAGCTCGAATTTCTTCC

### Protein expression and purification

PTPN2, PTPN2 C216G, and PTP1B were cloned into pGEX6P-1 expression vector and expressed in BL21 (DE3). When bacteria reached OD600 about 0.6, protein was then induced with 0.5 mM isopropyl-1-thio-B-D-galactopyranoside (IPTG) for 12 h at 25 °C. Cells were harvested and re-suspended in lysis buffer (50 mM Tris-HCl, pH7.5, 500 mM NaCl, 10 mM Imidazole, 10% Glycerol), lysed by high pressure crushing (ATS Engineering Limited, 800 bar, 15 min), and centrifugated (Beckman, 18000 rpm, 30 min, twice) to remove the cell debris. The supernatant was passed through a 5 mL Ni-column (GE) pre-equilibrated with lysis buffer. The His-GST-PTPN2, His-GST-PTPN2 C216G, or His-GST-PTP1B protein was eluted with 300 mM imidazole, then diluted with buffer (50 mM Tris-HCl, pH 7.5, 500 mM NaCl and 10% glycerol) to remove the imidazole. Subsequently, prescission protease was added to cut the N-terminal 6xHis-GST-Tag overnight at 4 °C. The target protein mixed with precision protease and tag was then concentrated with an Amicon Ultra-15 centrifugal filter unit (10,000 MWCO) (EMD Millipore) at 4 ℃. The target protein mixed with sumo protease and tag were then passed through a second 1 mL Ni-column and the flow-through solution containing target protein was collected and concentrated with an Amicon Ultra-15 centrifugal filter unit (10,000 MWCO) (EMD Millipore) at 4℃. Lastly, 1 ml protein was loaded onto the analytical size exclusion column Superdex 200 increase 10/300 GL (GE Healthcare) in running buffer (25 mM Tris-HCl, pH 7.5, 500 mM NaCl). All fractions were analyzed by SDS PAGE.

### Clinical samples

Eight blood samples from AML patients were obtained from Jiangsu Province Hospital of Chinese Medicine with patients’ informed consent. The human samples used in this study were approved by Ethics Committee of Jiangsu Province Hospital of Chinese Medicine. Clinicopathologic information of these eight patients was listed in Supplementary Table 1.

### Extraction of AML primary cells

Transfer 10 ml of whole blood into a 50 ml centrifuge tube, dilute with 10 ml of PBS solution and mix gently. Take two 15 ml centrifuge tubes and add 5 ml of Ficoll solution first. The diluted blood was then gently added to the upper layer of ficoll in both centrifuge tubes. Centrifuge at 2,000 rpm for 20 min. The cell layer where PBMC is located is white. At this point the layer can be pipetted into another clean 15 ml centrifuge tube. Add PBS to 10-15 ml, centrifuge at 1,500 rpm for 10 min and remove the supernatant, then add the medium with cytokines for the same operation. Add 5-10 ml of medium with cytokines to resuspend the cells for subsequent cell cultures or plates.

### Detection of phosphatase activity

The reaction system consisted of 50 mM Tris-Cl, PH = 7.2, 10 mM NaCl, 10% glycerol, and 0.1% bovine serum albumin (BSA). The reaction was carried out in 96-well plates, and the total reaction system was 100 μL, divided into three groups: experimental group, negative control group, and positive control group. In the experimental group, 48 μL of PTPN2 or PTP1B (manually purified) and 2 μL of compound (dissolved in DMSO) were added to each well, then mixed for 30 s and incubated at 37 ℃ for 15 min. Then 50 μL of 50 nM PNPP was added and mixed for 30 s and incubated at 37 ℃ for 5 Min, 100 μL 1M sodium hydroxide was added, and the absorbance at 405 nm was measured by a microplate reader, and the IC_50_ values of the target compounds against PTPN2 and PTP1B were calculated.

### Cell culture

The cell lines used in the experiment were all from the cell bank of Fu Heng Biology. Human acute myeloid leukemia cell lines MOLM13, NB4, K562, Jurkat and HL-60 were cultured in RPMI1640 medium containing 10% fetal bovine serum, while human bone marrow stromal cells HS-5 were grown in DMEM medium containing 10% fetal bovine serum. The cells were cultured in a 37 ℃, 5% CO_2_ cell incubator. All cells were tested annually for mycoplasma contamination using a PCR Mycoplasma detection kit and to prevent potential contamination, all media were supplemented with penicillin-streptomycin and mycoplasma scavicator according to the manufacturer’s instructions.

### Anti proliferation activity assay

The CCK8 kit was used to evaluate the antiproliferative activity of the compounds. Cells were incubated in a 37 ℃, 5% CO_2_ thermostatic cell incubator. The cells were seeded in 96-well plates with 100 μL of culture medium containing 8% fetal bovine serum at a density of 8000 cells/well. After 24 h of culture, the cells were treated with compound solution diluted in medium gradient (DMSO solubilization) for 72 hours, and then 10 μL of CCK8 reagent for 4 hours. The absorbance value of each well at 450 nm was detected with a Synergy H1 microplate reader (BioTek), and the results were analyzed using GraphPad Prism 8 to calculate the IC_50_ value.

### Cell growth assay

5000 tumor cells or shPTPN2 cells per well were seeded into 96-well plates and cultured for 4 days. CCK8 reagent was added daily to the Wells of each 96-well plate. The treated 96-well plates were incubated in an incubator for 4 hours. Absorbance was then measured at 450 nm wavelength using a microplate reader, and the results were analyzed using GraphPad Prism 8 to calculate IC_50_ values.

### Cell cycle assay

Cells were seeded into six-well plates, treated with test compounds for 24 h, then washed twice with PBS, removed, added with 70% ethanol, gently beaten, and fixed at 4 °C for 12-24 h. Centrifugation at 1000g for 5 min, carefully aspirate the supernatant of the precipitated cells, add 1 mL PBS, resuspended in 0.5 mL Krishan’s buffer, supplemented with 0.05 mg/mL PI, 0.1% trisodium citrate, The cells were incubated at 37 °C for 30 min with 0.02 mg/mL ribonuclease A and 0.3% NP-40, and then directly applied to flow cytometry (BD, USA) and analyzed with FlowJo V10 software.

### Apoptosis assay

Cells were seeded into six-well plates and treated with test compounds for 24 hours. Cells were then washed twice with cold PBS and re-suspended in binding buffer (0.1 M HEPES (pH = 7.4), 1.4 M NaCl, 25 mM CaCl_2_). A 100 μL volume of solution was transferred to a 5 mL flow tube. 5 μL Annexin V-APC and 5 μL PI were added to each tube. The cell suspension was gently vortexed and incubated at room temperature in darkness for 30 min. Apoptosis was determined by flow cytometry (BD, USA) at 633 nm excitation, and the results were analyzed by FlowJo V10 software.

### Real-time quantitative PCR

Total RNA isolated from cells with RNA-Easy reagent was reverse transcribed using the Hiscript 1^st^ Strand cDNA Synthesis kit. RT-qPCR reactions were performed with ChamQ SYBR qPCR Master Mix. Relative gene expression was calculated by relative quantification method. Human PTPN2 primers: Forward (5’-3’): ATCGAGCGGGAGTTCGA, Reverse (5’-3’): TCTGGAAACTTGGCCACTC; Human ACTIN primers: Forward (5’-3’): GTGGCCGAGGACTTTGATTG, Reverse (5’-3’): TGGAVTTGGGAGAGGCTGG.

### Lentivirus production and infection

According to the gene sequence of PTPN2, the target sequence of RNAi was designed, and the target sequence was constructed into the corresponding lentiviral vector, which was then transformed, identified by colony PCR, sequenced, and then extracted for transfection into HEK-293T cells to obtain the virus. The lentivirus was directly infected with MOLM13 cells. MOLM13 cell lines containing lentivirus were infected at 32 ℃ and 1200 RPM for 90 minutes and cultured continuously. After 48 hours of infection, 2 μg/mL puromycin was added to MOLM13 cells, and finally the successfully infected MOLM13 cells were selected. PTPN2 shRNA sequences: CCGGGATGACCAAGAGATGCTGTTTCTCGAGAAACAGCATCTCTTGGTCATCTTTTTG.

### RNA-Seq sequencing

Total RNA was extracted from PTPN2 knock down cells using the Trizol kit (Invitrogen, Carlsbad, CA, USA). RNA quality was assessed using an Agilent 2100 bioanalyzer (Agilent Technologies, Palo Alto, CA, USA) and checked by electrophoresis on RNase-free agarose gels. After total RNA extraction, eukaryotic mRNA was enriched with Oligo (dT) beads and prokaryotic mRNA was enriched by removing rRNA with Ribo-ZerotM Magnetic Kit (Epicentre, Madison, WI, USA). The enriched mRNA fragments were then split into short fragments using lysates and reverse transcribed into a strand of cDNA using random primers. The second strand of cDNA is synthesized by DNA polymerase I RNase H dNTP and buffer. The cDNA fragment was then purified with QiaQuick PCR extraction Kit (Qiagen, Venlo, Netherlands), the end was repaired, the A base was added, and ligated to an Illumina sequencing adapter, The ligation products were screened by agarose gel electrophoresis, amplified by PCR, and sequenced by Illumina Novaseq 6000.

### High throughput protein phosphorylation modification omics

Protein was extracted from each sample and the same amount was taken for enzymatic hydrolysis. The final concentration of 20% TCA was slowly added, and the mixture was vortexed and precipitated for 2h at 4 ℃. Centrifugation at 4500 g for 5min, discard the supernatant, and wash the precipitate with pre-cooled acetone for 2-3 times. After the precipitation was dried, the final concentration of 200 mM TEAB was added, the precipitation was broken up by ultrasound, trypsin was added, and the precipitation was enzymolized overnight. Dithiothreitol (DTT) was added to make the final concentration of 5 mM and then reduced at 56 ℃ for 30 min. Then iodoacetamide (IAA) was added to make the final concentration of 11 mM, and the mixture was incubated for 15 min at room temperature under light. The peptides were solubilized with 0.5 M TEAB and labeled according to the instructions of TMT kit. Peptides were fractionated by high-pH reverse-phase HPLC. The peptides were dissolved in enrichment buffer solution (50% acetonitrile /0.5% acetic acid), and the supernatant was transferred to pre-washed IMAC material and incubated with gentle shaking on a rotary shaker. At the end of incubation, the material was washed three times with buffer solution 50% acetonitrile /0.5% acetic acid and 30% acetonitrile /0.1% trifluoroacetic acid. Finally, the phosphopeptides were eluted with 10% ammonia, and the eluent was collected and vacuumed, and then the eluent was vacuumed and dried, and then the phosphopeptides were used for liquid mass spectrometry analysis. The peptides were dissolved in liquid chromatography mobile phase A and separated by EASY-nLC 1200 ultra-high performance liquid system. The peptides were separated by a UHPLC system and then injected into an NSI ion source for ionization and then analyzed by Q Exactive™ HF-X mass spectrometry.

### MST assays

Microthermophoresis Microscale thermophoresis (MST) is a technique for analyzing biomolecular interactions. The technique is based on the thermophoresis of biomolecules. Nanotemper microscale thermophoresis uses an infrared laser for localized heating leading to directional movement of molecules, followed by fluorescence analysis of molecular distribution ratios in the field of temperature gradients. The affinity of PTPN2 and PTPN2 C216G for K73 was examined in vitro using micro-thermophoresis. The affinity between PTPN2 or PTPN2 C216G and K73 was measured by observing the changes in the conformational size, charge, and solvation state of the target molecule after binding of PTPN2 or PTPN2 C216G to K73. Purified recombinant PTPN2 protein (20 μM) was labeled according to the protein labeling kit RED-NHS (Nanotemper, Cat. No. L001). The K73 compound was diluted into 16 concentration gradients with 1xPBS-P using the fold-and-half dilution method. The labeled PTPN2 protein or PTPN2 C216G protein was mixed with different concentrations of K73 in equal volumes and incubated at room temperature for 30 min. MST data were then collected at 20% IR laser power and 20% light emitting diode power. The data were analyzed and K_d_ was determined using Nanotemper analysis software.

### SPR assays

Biacore S200 (General Electric Company, USA) was used to perform the SPR binding experiment. K73 was dissolved in PBS-P buffer. After diluting the PTPN2 protein with sodium acetate solution (pH = 4.5) to 20 μg/mL, the protein was conjugated to CM5 chip. K73 was injected at 25 ℃ at a flow rate of 0.5 μL/s at various concentration. The Langmuir binding model was utilized in order to compute the K_d_ value.

### Western blotting analysis

For MG132 experiment, cells were treated with 5 mM MG132 for 12 h, and cells were collected for western blot analysis. Cells were treated with vehicle or the specified K73 concentration for 72 h, and treated cells were harvested and lysed by sonication in receptor lysis buffer (RLB), phosphatase inhibitors, and protease inhibitor mix. Lysates from cells and tumor tissues were quantitated and 20 to 50 μg of protein lysates were boiled in an SDS sample buffer, size fractionated by SDS-PAGE, and transferred onto a PVDF membrane (Immobilon). After blocking in 5% nonfat dry milk (or 3% BSA), membranes were incubated with the following primary antibodies overnight: PTPN2 rabbit polyclonal antibody (Proteintech, Cat#: 11214-1-AP, 1:1000), RB rabbit polyclonal antibody (Proteintech, Cat#: 10048-2-Ig, 1:5000), Phospho-Rb (Ser807/811) rabbit monoclonal antibody (Cell Signaling Technology, Cat#: 8516, 1:1000), CDK4 rabbit monoclonal antibody (Cell Signaling Technology, Cat#: 12790, 1:1000), CDK6 mouse monoclonal antibody (Cell Signaling Technology, Cat#: 3136, 1:2000), P21 rabbit polyclonal antibody (Proteintech, Cat#: 10355-1-AP, 1:1000), P27 rabbit monoclonal antibody (Cell Signaling Technology, Cat#: 3686, 1:1000), P53 rabbit monoclonal antibody (Cell Signaling Technology, Cat#: 2527, 1:1000), XIAP rabbit polyclonal antibody (Proteintech, Cat#: 10037-1-Ig, 1:1000), C-MYC rabbit polyclonal antibody (Proteintech, Cat#: 10828-1-AP, 1:2000), C-MYC (phospho T58) rabbit polyclonal antibody (abcam, Cat#: ab28842, 1:1000), BCL-2 rabbit polyclonal antibody (Proteintech, Cat#: 12789-1-AP, 1:2000), STAT1 rabbit monoclonal antibody (Cell Signaling Technology, Cat#: 9172, 1:1000), Phospho-STAT1 (Tyr701) rabbit monoclonal antibody (Cell Signaling Technology, Cat#: 9167, 1:1000), Vinculin rabbit polyclonal antibody (Proteintech, Cat#: 26520-1-AP, 1:1000), Alpha Tubulin mouse monoclonal antibody (Proteintech, Cat#: 66031-1-Ig, 1:20000), GAPDH rabbit monoclonal antibody (Cell Signaling Technology, Cat#: 2118, 1:1000). Following three washes in PBST, the blots were incubated with secondary antibody Goat Anti-Mouse IgG, H&L Chain Specific Peroxidase Conjugate (Merck, Cat#: 401215-2 ML, 1:5000), or Goat Anti-Rabbit IgG, H & L Chain Specific Peroxidase Conjugate (Merck, Cat#: 401315-2 ML, 1:5000). Proteins were detected by chemiluminscent detection system (Tanon, Shanghai, China) and analyzed by Image J software.

### Transient transfection

All plasmids including PTPN2, PTPN2 C216G, C-MYC and C-MYC T58A were constructed. When cells grew to logarithmic phase, the cells were spread in 6-well plates with 500,000 cells per well, transiently transfected using polyethyleneimine (PEI), and transfection complexes were prepared according to PEI: DNA = 3:1 and the medium was changed after 18 h of staining, and the cells were collected 72 h later.

### GST-C-MYC recombinant protein

MOLM13 cells were selected as the over-expression cell line. After MOLM13 cells reached at logarithmic stage, 9 mL RPMI1640 cell suspension containing 10%FBS was prepared, 4×10^6^ cells were counted, and 10 cm dishes were spread. The cells were divided into Vector-pCDNA3.1(+) group and GST-C-MYC-pCDNA3.1(+) group. 1 mL of transfection complex was prepared according to PEI: DNA=3:1. The transfection complex was added drop by drop to the culture dish, and the solution was changed 18 h after transfection, and the cells were collected 72 h later. Some cells were collected from each group and over-expression was verified by western blot. The remaining cells in GST-C-MYC-pCDNA3.1(+) group were lysed with western and IP lysates.

### GST pull down assay

Recombinant GST protein or GST-PTPN2 protein immobilized on glutathione Sepharose 4B beads (GE) in a binding buffer containing 25 mM HEPES, pH 7.5, 200 mM NaCl, and 1 mM EDTA for 4 hr incubation at 4 ℃. Whole cell lysates were prepared in mild lysis buffer containing 50 mM Tris-HCl, pH 7.6, 150 mM NaCl, 1% NP-40, 0.5% sodium deoxycholate and 0.1% SDS, supplemented with 1 mM PMSF, protease and phosphatase inhibitor cocktails on ice for 30 min. Followed by ultrasonic crushing, the whole cell lysates were centrifugated at 12,500 rpm for 15 min. The supernatant was added to recombinant GST protein or GST-PTPN2 protein for overnight incubation at 4 ℃. The beads were washed three times with binding buffer and eluted with the same buffer plus 10 mM Glutathione (adjusted to pH 7.5). The eluted proteins were visualized with Western Blot.

### Co-IP assay

Protein lysates were prepared from MOLM13 cells in an Enhanced Lysis Buffer and a mixture of protease inhibitors by using a Co-IP kit (ACE Biotechnology). Followed by ultrasonic crushing, the whole cell lysates were centrifugated at 12,500 rpm for 15 min. The supernatant was added to 20 μL of Protein A/G MagPoly Beads in 1 mL Enhanced Lysis Buffer. The lysates were then incubated with 4 μg anti-PTPN2 antibody or control IgG overnight at 4 ℃. The complexes were washed three times with Enhanced Lysis Buffer and eluted from the beads with IP Elution Buffer and Neutralization Buffer. The eluted proteins were detected by Western Blot.

### Immunoprecipitation

The cells were lysed using a lysis buffer and placed on ice for a duration of 5 minutes. To release the nuclear proteins, the lysates were subjected to centrifugation at 13000g for 10 minutes after thorough mixing. For the subsequent steps, 10 μl of anti-PTPN2 antibody was added to the proteins and allowed to incubate for a period of 2 hours at a temperature of 4 °C. Additionally, rabbit IgG was also added to the proteins and allowed to incubate for 2 hours, under the same temperature conditions. Overnight co-incubation at 4 °C was carried out with 2% BSA, antibodies, and 10 mg of protein A-Sepharose beads. The proteins were then eluted thrice, and the eluate was fractionated through SDS-PAGE. Subsequently, the gel pieces were incubated in the dark for 45 minutes at room temperature. Washing of the gel pieces was performed using 50 mM NH_4_HCO_3_, followed by dehydration using 100% acetonitrile. The dehydrated gel pieces were then resuspended in 50 mM NH_4_HCO_3_ and trypsin for 1 hour on ice. After removing the excess liquid, trypsin digestion was carried out at 37 °C overnight. Further extraction of peptides was performed using 50% acetonitrile/5% formic acid and followed by 100% acetonitrile. The extracted peptides were dried completely and resuspended in 2% acetonitrile/0.1% formic acid. The tryptic peptides were dissolved in solvent A (0.1% formic acid) and directly loaded onto a reversed-phase analytical column created in-house. MS/MS analysis, utilizing Q ExactiveTM Plus (Thermo) coupled online to UPLC, was conducted subsequent to NSI source application. The full scan was performed within an m/z range of 350 to 1800, and the intact peptides were detected at a resolution of 70,000 in the Orbitrap. Peptides were selected for MS/MS analysis, and the resulting data were processed using Proteome Discoverer 1.3.

### In vitro dephosphorylation assay

C-MYC was previously overexpressed in MOLM13 cells and purified to obtain C-MYC. Post-translational modification occurred in eukaryotic cells, and the expressed and purified C-MYC was phosphorylated at threonine 58. Naturally modified C-MYC was used as a substrate, and the phosphatases were recombinant proteins PTPN2 and PTPN2 C216G. Dephosphorylation buffer (20 mM Tris-HCl (pH 7.5), 100 mM NaCl, 0.57 mM EDTA, 0.033% BSA, and 1 mM DTT) was used. p-C-MYC (T58), different concentrations of PTPN2 (0.25 μM, 0.5 μM,1 μM) and mutant PTPN2 C216G (1 μM) were incubated at 30 °C for 1 h, and 4 x Loading Buffer was added to prepare samples. The levels of p-C-MYC (T58) were verified by western blot analysis. p-C-MYC (T58), 1 μM of PTPN2 and different concentrations of K73 were incubated at 30 ℃ for 1 h, and 4 x Loading Buffer was added to prepare samples. The levels of p-C-MYC (T58) were verified by western blot.

### Acute toxicity studies

ICR mice were used for acute toxicity tests to evaluate the safety of the selected compounds. Four dose groups were set up, and the calculated dose was given once for two weeks. The body weight of the animals was measured every two days, and the health status of the animals was recorded every day, including the appearance characteristics, activity status, mental state, food and water intake, and death. After 14 days of dosing, the animals were euthanized. The heart, liver, and spleen were collected. Histological evaluation of lungs and kidneys was performed, and the body weight curve of the animals was made and analyzed.

### In vivo anti-tumor activity

To establish the shNC group (n=5) and shPTPN2 group (n=5), BALB/c nude mice were administered a subcutaneous injection of 1 × 10^7^ MOLM13 cells harboring shNC and shPTPN2, respectively. Tumor volumes and body weight measurements were taken at two-day intervals once tumors had formed. At 16 days, the mice from both groups were euthanized for ethical reasons, and tumor tissues were weighed and photographed.

MOLM13-Luciferase (5 × 10^5^) tumor cells were injected into the tail vein of M-NSG mice to establish MOLM13-Luciferase transplantation tumor model. On the second day after inoculation, tumor-bearing mice were randomly divided into two groups according to Flux value: control group and compound K73 group (10 mg/kg), with 5 mice in each group. On the 3rd day after inoculation, the mice were administered once a day for 10 days. The body weight of mice, the changes of tumor fluorescence value, and the death of mice were recorded every day.

### Statistics and reproducibility

In order to compare the difference between two groups, the unpaired, a two-tailed Student’s t-test was employed using the SPSS 22.0 software, unless otherwise specified in the figure captions. To conduct bioinformatics analysis, the Student’s t-test and Mann-Whitney Wilcoxon test were utilized to compare continuous variables between two groups. For the assessment of variance among multiple groups, the analysis of variance (ANOVA) was employed. Relapse-free survival analysis was performed using Kaplan-Meier plots and log-rank tests. Mean ± SD values were used to express the results, and the number of replicates is specified in the figure captions. A statistically significant difference was considered when P < 0.05.

## Data availability

The raw RNA-seq data generated in this study have been deposited in the Genome Sequence Archive for Human (https://ngdc.cncb.ac.cn/gsa-human/) under the accession number: HRA006552. The mass spectrometry proteomics as well as identified and significantly regulated phosphosites data have been deposited to the ProteomeXchange Consortium (https://proteomecentral.proteomexchange.org) via the iProX partner repository with the dataset identifier PXD048854.

## Acknowledgments

This study was supported by the National Key R&D Program of China (2022YFA1303803), National Natural Science Foundation of China (82073701, 82373738, 82304296), Natural Science Foundation of Jiangsu Province (BK20231013) and the Project Program of State Key Laboratory of Natural Medicines, China Pharmaceutical University (SKLNMZZ202209). This study was also supported by the Fundamental Research Funds for the Central Universities (2632023GR22).

## Author contributions

W.K., J.J., X.W., and D.W. conceived the project, which had leadership from W.G., H.H., Y.X., and P.Y. All authors contributed to manuscript writing, review, and editing. W.K., D.W., X.W., and J.D. contributed to cancer database analysis. W.K., J.J., D.W., X.W., M.J., J.D. and F.H. contributed to biological experiments, including Protein expression and purification, cell proliferation assays, lentivirus production and infection, real-time quantitative PCR, immunoprecipitation, western blotting, colony formation assays, migration and invasion assays, cell cycle and apoptosis assays, RNA-seq sequencing, GST pull down etc. W.K., W.W. and W.M. contributed to structure-based virtual docking. W.K., Y.Z., K.Y., W.W., and P.Y. contributed to the design and synthesis of PTPN2 inhibitors. W.K., D.W., and P.Y. contributed to in vivo acute toxicity properties and anti-tumor activity evaluation in vitro and in vivo. Project administration and funding acquisition: W.K. and P.Y. All authors contributed to data analysis and agreed to the published version of the manuscript.

